# Decreased tRNA abundance contributes to decreased translation elongation rate in a prolonged mitosis

**DOI:** 10.64898/2026.01.13.699274

**Authors:** Emily Sparago, Ryan Johnston, Shawn M. Lyons, Michael D. Blower

## Abstract

Cells undergo dramatic structural rearrangements upon entering mitosis. In addition, the biochemistry of mitotic cells is dramatically altered, including a significant decrease in protein synthesis. The majority of studies of mitotic translation have used cells synchronized by cell cycle altering drugs in transformed cells and much less is known about mitotic translation in primary cells under native conditions. Previous work has found that mitosis activates the integrated stress response (ISR) to trigger eIF2α phosphorylation, but little is known about the input for this response. In this study, we focus on mitotic translational regulation in an immortalized, non-transformed cell line. We confirm decreased mitotic protein synthesis in primary cells and under native conditions. Additionally, we confirm activation of the ISR by phosphorylation of eIF2α during both normal and prolonged mitosis. Interestingly, we also find that decreased translational elongation during mitosis, as evidenced by increased eEF2 phosphorylation and a slower elongation rate. Analysis of mitotic ribosome profiling data revealed an increase in pausing at Alanine-GCG codons during mitosis and a decreased abundance of its cognate tRNA-Ala^CGC^ by northern blotting. Decreased tRNA-Ala^CGC^ is likely sustained by the inability to synthesize additional tRNA due to RNAPol III inhibition in mitosis, yielding an stronger effect with an increased time in mitosis. These results suggest that decreased translation elongation in mitosis triggers inhibition of initiation to decrease global protein synthesis.

## Introduction

Mitosis leads to a major reorganization of nearly every structure in the cell. In addition, cellular biochemistry is altered to promote cell division. For example, the vast majority of transcription is repressed during mitosis in mammalian cells. RNA Polymerase II (RNAPol II)-dependent transcription is suppressed by phosphorylation of key transcription initiation factors and leads to dissociation of polymerases from chromatin (1–3). Studies confirm that transcription by the three main RNA polymerases, Pol I, Pol II, and Pol III are inhibited during mitosis, which primarily transcribe rRNA, mRNA, and tRNA, respectively (4, 5).

Additionally, translation is attenuated during mitosis (4, 6–13). Translation decreases during interphase under various conditions of stress, which can be mediated at initiation or elongation. Activation of the integrated stress response (ISR) is characterized by the phosphorylation of eIF2α at serine 51, preventing the exchange of GDP to GTP to form the ternary complex to prevent ribosomes from transitioning from initiation to elongation (14). The four main kinases which phosphorylate eIF2α are double-stranded RNA-dependent protein kinase (PKR), PKR-like endoplasmic reticulum kinase (PERK), heme-regulated eIF2α kinase (HRI), and general control nonderepressible 2 (GCN2). These kinases respond to distinct stimuli; PKR recognizes double-stranded RNA, such as that produced in viral infection (15), while PERK is activated through ER stress (16). HRI is predominately expressed in erythropoietic cells and recognizes heme availability; however, this kinase also responds to oxidative stress produced by arsenite (17). Finally, GCN2 is activated by amino acid deprivation, depletion of tRNAs, or ribosomal stalling and collisions (18, 19). Alternatively, translation initiation can be inhibited through binding of 4E-BP to the canonical cap-binding protein, eIF4E1, which prevents the recruitment of the other proteins in the eIF4F complex. 4E-BP binding to eIF4E is triggered by inhibition of the mTORC1 pathway, which is normally activated by recognition of certain cell proliferation-related signals, such as growth factors, amino acids, energy and oxygen (20).

In addition to initiation, translation can be regulated through decreased elongation which primarily focuses on the elongation factor, eEF2. eEF2 promotes ribosome translocation, moving the peptidyl-tRNA from the ribosomal aminoacyl (A-) site to peptidyl (P-) site, but its function is halted during stress, such as amino acid starvation, to prevent an empty A site (21). Previous research found that mitosis exhibited both a decreased elongation rate and decrease in translation initiation (22–25). Causes for an empty A site include amino acid starvation, leading to a decrease in charged tRNAs, or a depletion of total tRNA abundance (26). Growing research has demonstrated dramatic dynamics of tRNA abundance, including cleavage and degradation, in response to stress, cancer, and DNA damage (27–30). Any of these changes could ultimately lead to an empty A site, triggering decreased translational elongation.

Previous studies of translation during mitosis employ a range of methods to enrich a mitotic population, utilizing various drugs to synchronize cells for an extended mitosis. In addition, previous work on mitotic translation often focuses on cancer cells, which have modified mechanisms to bypass typical cell cycle checkpoints (31). In this work, we explore the translation regulation of a normal, immortalized primary cell line, RPE-1 cells. We find a global decrease of translation in mitosis, with regulation at both translation initiation and elongation. Interestingly, we identify mitosis-specific pausing at Ala-GCG codons, which is triggered by a decrease in tRNA availability. We propose that lack of a specific tRNA triggers phosphorylation of eEF2 to slow translation elongation and ultimately leads to phosphorylation eIF2α to slow translation initiation. We speculate that the lack of tRNA is likely caused by increased degradation of tRNA-Ala^CGC^ and is sustained by inhibition of global transcription, including RNAPolIII, failing to replenish tRNAs for effective translation. As a result, translational repression increases in severity with increased time in mitosis.

## Results

### Translation is suppressed in native and drug-arrested mitotic cells

Previous work on mitotic translation covers a range of cell types, though often focusing on cancer cells which have adapted unique mechanisms to bypass typical cell cycle checkpoints (31). Many studies have used a mitotic population enriched by 16-24-hour cell synchronization (4, 6, 7, 9–11, 13, 25, 32, 33). Prior research has explored key translational differences in mitosis in cancer and primary cells (34). Here we examined whether differences exist between cells that go through mitosis for a typical length of time (cycling mitosis) and those synchronized for 12-16 hours (prolonged mitosis) in a single, immortalized, non-transformed cell type, hTERT-RPE1 cells.

It has been shown that eIF2α is phosphorylated during a prolonged mitosis, indicating a stress response (24, 25). We confirmed our RPE-1 cells indeed respond to stress through treatment with sodium arsenite (AsO_2_), which is known to decrease protein synthesis through eIF2α phosphorylation (35, 17). Using polysome profiling, we observed a decrease in polysome:monosome ratio (PM) characterized by the increase in the 80S peak and subsequent loss of polysomes (Figure 1A, E), which reflects translation efficiency. Our cycling mitotic population uses an enrichment from cells previously synchronized in G2 with RO-3306 and subsequently collected by mitotic shake-off (Supplemental Figure 1). Cycling mitotic cells exhibit a clear decrease in translation relative to interphase cells (Figure 1B, E). Strikingly, the samples synchronized in mitosis overnight by Nocodazole or STLC treatment show a more severe decrease in translation efficiency, characterized by a major increase in the monosome peak and loss of polysomes (Figure 1C, E). Comparison of cycling and prolonged mitotic cells revealed that translation efficiency is reduced the longer cells are arrested in mitosis, as evidenced by the difference in the 80S peak (Figure 1D, E). We confirmed decreased mitotic translation in Hap1 cells within an asynchronous population, void of drug treatment, through puromycin immunofluorescence; mitotic cells show less puromycin incorporation, supporting a decrease in protein synthesis during mitosis irrespective of drug treatment (Supplemental Figure 2).

**Figure 1.**
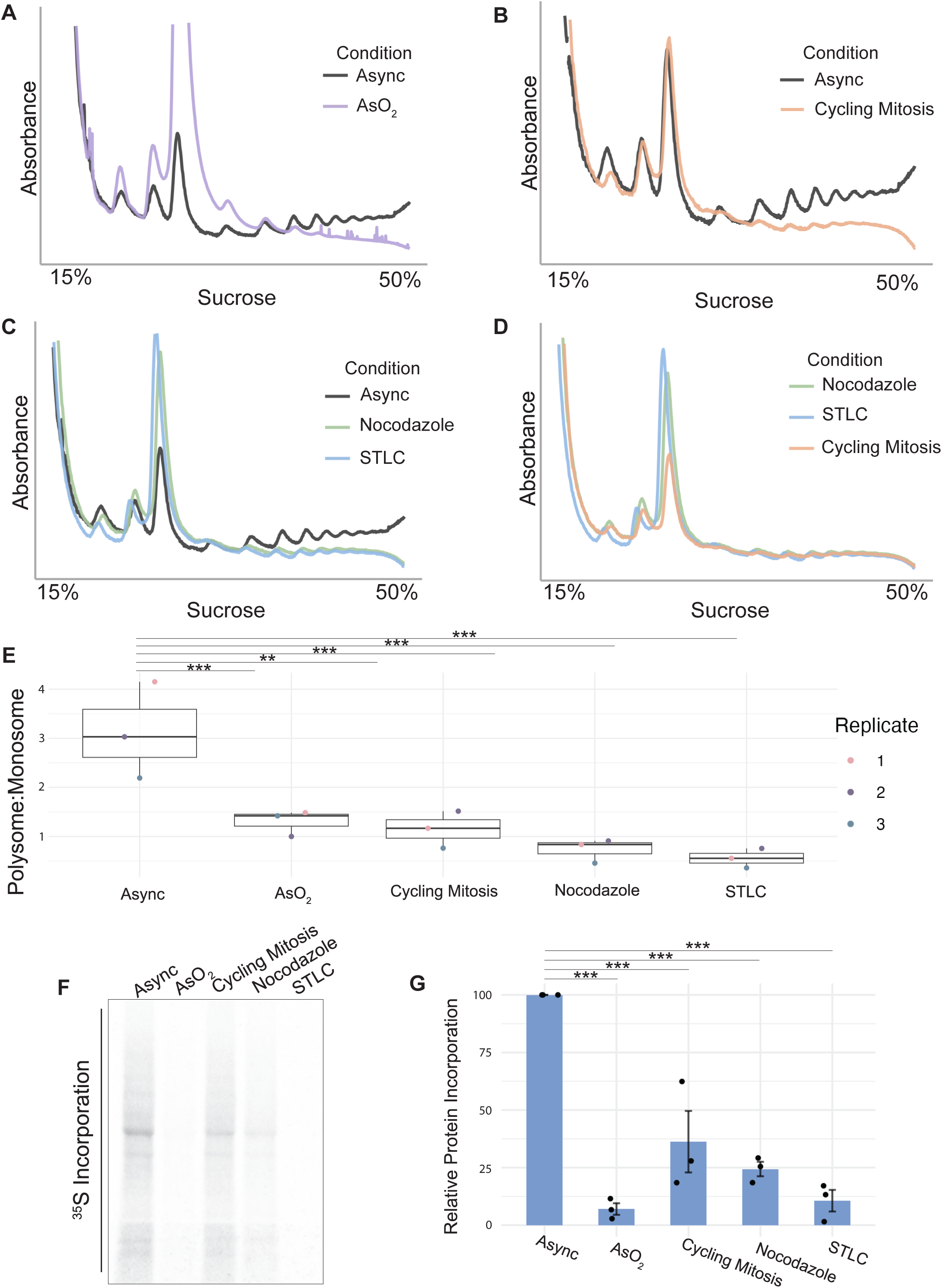
Basal translation is decreased in mitosis. Representative polysome trace of RPE-1 asynchronous cells compared to **(A)** sodium arsenite (NaAsO_2_) treated; **(B)** cycling mitotic cells by an RO-3306 release; **(C)** mitotic cells enriched by Nocodazole and STLC treated cells (16h); **(D)** and all mitotic-enriched samples. **(E)** Relative translation efficiency by polysome to monosome ratio. Significance determined by one-way ANOVA with Dunnett’s post-hoc test, asynchronous as reference. **(F, G)** Metabolic labeling by ^35^S shows relative protein synthesis. Significance determined by one-way ANOVA with a Dunnett’s post-hoc test, asynchronous as reference.

To confirm this translational effect using an orthogonal approach, we performed metabolic labeling with [^35^S]-methionine/cysteine. Our positive control, AsO_2_, shows a significant decrease in protein synthesis. Similarly, the mitotic populations, both cycling and prolonged, show a comparable decrease in protein synthesis (Figure 1F, G). These combined results are consistent with previous work (7, 8). Decreased mitotic translation suggests mitosis to be a stress to cells (36).

### Bypass of eIF2α phosphorylation does not rescue translation in mitosis

The ISR is a critical pathway that regulates protein synthesis upon exposure to stress and is initiated by the phosphorylation of eIF2α to prevent translation initiation (14). To determine if mitosis triggers the ISR, we analyzed eIF2α phosphorylation in asynchronous, mitotic, and AsO_2_-treated cells. We observed a significant increase in the levels of phosphorylated eIF2α in all mitotic and AsO_2_-treated samples, relative to asynchronous, by comparison of phosphorylated to total eIF2α intensity (Figure 2A, B). This indicates translation initiation is decreased in mitosis consistent with previous work showing that cells begin to regulate translation at the G2/M boundary through eIF2α phosphorylation ((25), Figure 2A, B).

**Figure 2.**
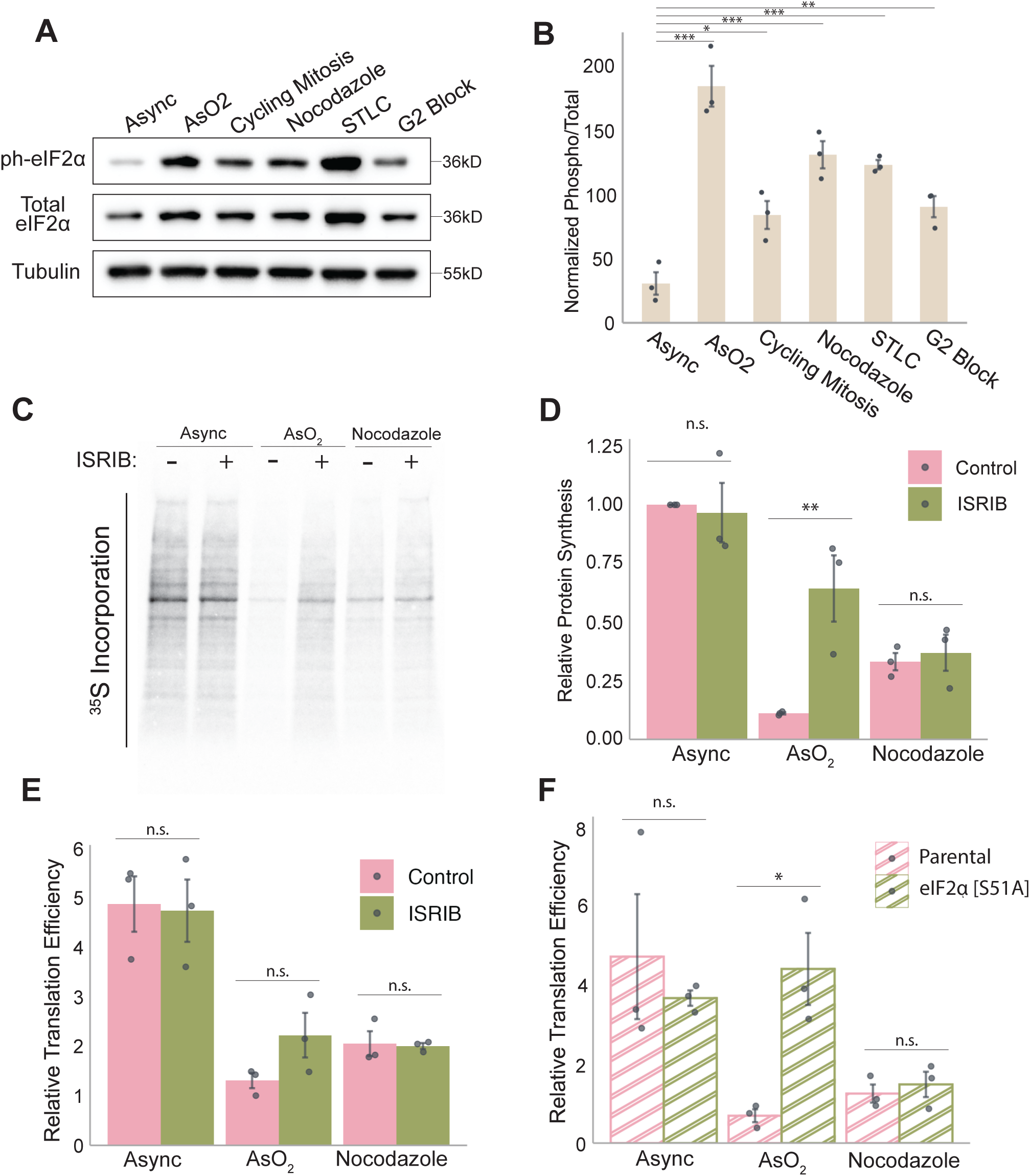
Contribution of eIF2α phosphorylation to mitotic translation inhibition. **(A)** Western blot of phosphorylated eIF2α and total eIF2α in RPE-1 cells in asynchronous and mitotic-enriched cells. **(B)** Quantification normalized to loading control, comparing phosphorylated to total eIF2α. Statistical significance determined by a one-way ANOVA with Dunnett’s post-hoc test, asynchronous as reference. Error bars represent the standard deviation, n=3 biological replicates. **(C)** ^35^S metabolic labeling for RPE-1 cells with ISRIB treatment in asynchronous and Nocodazole-enriched cells. **(D)** Quantification of metabolic labeling by two-way ANOVA with Tukey’s post-hoc test. Error bars represent the standard deviation, n=3 biological replicates. **(E)** Quantification of polysome analysis by polysome to monosome ratio for ISRIB-treated RPE-1 cells; statistical significance determined by two-way ANOVA with Tukey’s post-hoc test. Error bars represent the standard deviation, n=3 biological replicates. **(F)** Quantification of polysome analysis by polysome to monosome ratio of Hap-1 parental and eIF2α [S51A] cells. Statistical significance determined by two-way ANOVA with Tukey’s post-hoc test. Error bars represent the standard deviation, n=3 biological replicates.

To determine if eIF2α phosphorylation is the main mechanism responsible for the decrease in translation during mitosis, we blunted ISR-mediated translation inhibition by co-treatment with ISRIB, which allows eIF2α to exchange GDP to GTP irrespective of the phosphorylation state of eIF2α (37). We confirmed drug efficacy by metabolic labeling of cells treated with AsO_2_ alone or co-treated with ISRIB. ISRIB treatment resulted in significantly more protein synthesis in AsO_2_-treated cells. In contrast, protein synthesis remains decreased in mitotic cells with ISRIB treatment, relative to asynchronous cells (Figure 1C, D). To confirm this using an orthogonal method, we performed polysome profiling with the AsO_2_ and ISRIB treatment and found that ISRIB partially rescued translation in AsO_2_-treated cells. In contrast, there was no change in translation efficiency with or without ISRIB in nocodazole-treated mitotic cells (Figure 2E, S1). We further evaluated the effects of eIF2α phosphorylation by using eIF2α [S51A] Hap1 cells in which eIF2α cannot be phosphorylated (38). Polysome profiling reveals recovery of translation efficiency in eIF2α [S51A] Hap1 cells with AsO_2_ treatment, but no effect in a prolonged mitosis (Figure 2F). These data suggest that eIF2α is not the sole mechanism by which translation is decreased in a prolonged mitosis.

### 4E-BP is hyperphosphorylated in mitosis

Translation initiation can also be inhibited by dephosphorylation of 4E-BP1 following inhibition of mTOR signaling. Under ideal conditions, mTOR constitutively phosphorylates 4E-BP1, which maintains it in an inactive state. During stress, mTOR is inactivated, which results in 4E-BP1 dephosphorylation, allowing it to bind eIF4E. This prevents formation of eIF4F on the cap of the mRNA to initiate canonical, cap-dependent translation (39). To evaluate whether decreased eIF4F formation contributes to decreased translation efficiency in mitosis, we examined the phosphorylation of total 4E-BP1. We found hyperphosphorylation of 4E-BP1 in both cycling and prolonged mitosis (Nocodazole-treated). We also observed a major dephosphorylation of 4E-BP1 in a G2 block, consistent with previously reported literature ((25), Figure 3A). Additionally, we recognize a slight decrease in total 4E-BP1 in nocodazole-treated cells, suggesting that 4E-BP1 may be degraded during a prolonged mitosis (Figure 3A, B).

**Figure 3.**
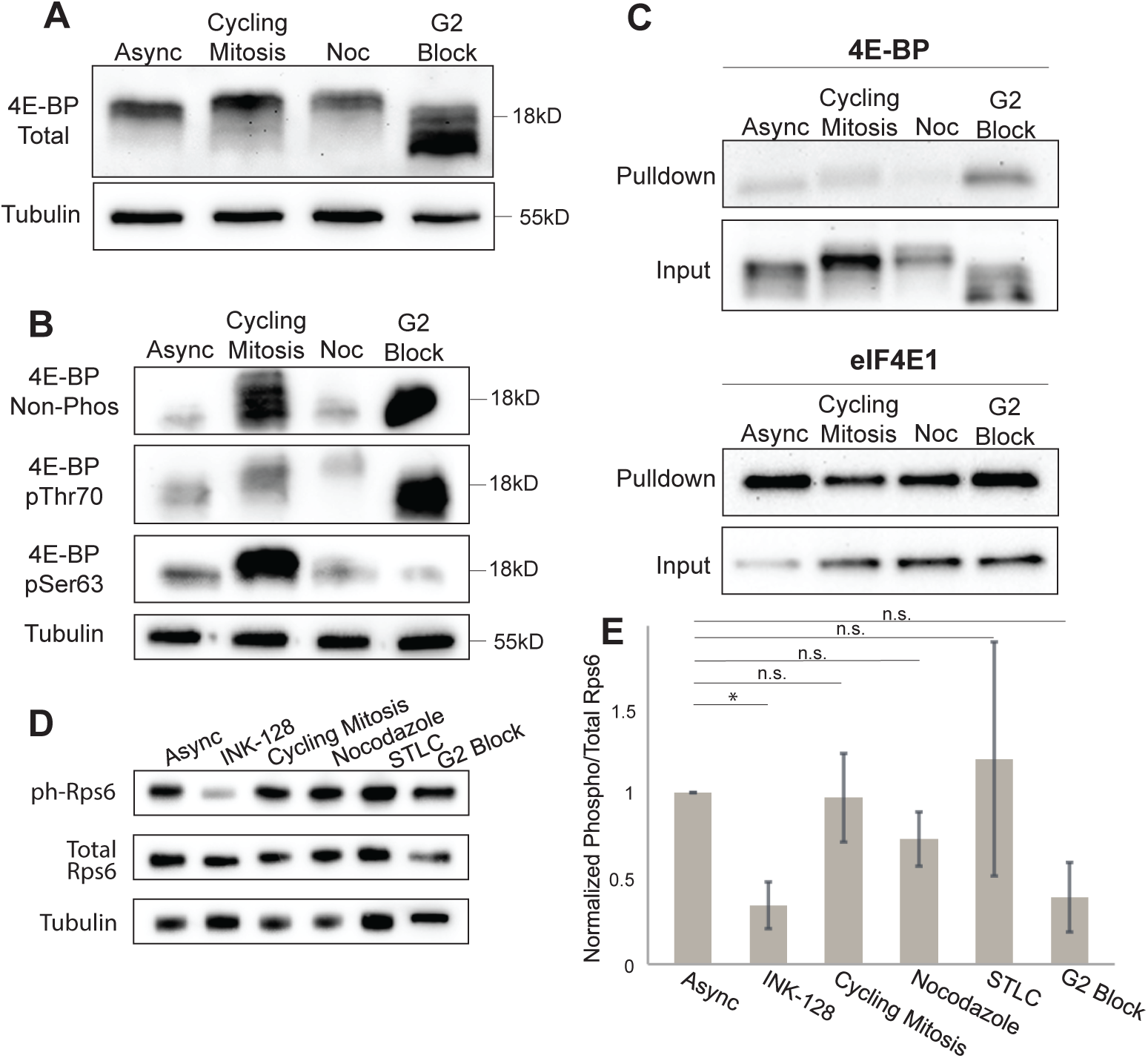
4E-BP is hyperphosphorylated in mitosis. **(A)** Western blot of total 4E-BP in RPE-1 asynchronous, mitotic-enriched, and G2-synchronized cells. **(B)** Western blot of non-phosphorylated, phosphorylated Threonine 70, and phosphorylated Serine 63 4E-BP. **(C)** Cap-pulldown analysis with m^7^GTP agarose beads with asynchronous, mitotic-enriched, and G2-synchronized RPE-1 cells. Western blot of 4E-BP total and eIF4E1. **(D)** Western blot analysis of phosphorylated and total Rps6 in RPE-1 asynchronous cells, mitotic-enriched cells, and treated with an mTOR inhibitor (INK-128). **(E)** Quantification of normalized phosphorylated to total Rps6. Significance determined by one-way ANOVA with Dunnett’s post-hoc, asynchronous reference. Error bars represent the standard deviation, n=3 biological replicates

To determine if 4E-BP1 shows increased binding to eIF4E1 during mitosis, we utilized m^7^GTP-sepharose beads in a cap-binding pulldown assay. If 4E-BP1 is bound to eIF4E1 on the mRNA cap, we would expect more 4E-BP1 to be pulled down with the m^7^GTP-sepharose beads, as in a G2 block (25). However, in its hyperphosphorylated state during mitosis, 4E-BP1 is not pulled down with eIF4E1 more than asynchronous cells (Figure 3C). Therefore, we conclude that 4E-BP1 binding to the eIF4E1 is not a mechanism by which translation is inhibited during mitosis.

To determine if mTOR is inactivated during mitosis, we analyzed phosphorylation of the mTOR substrate RPS6. We observed a slight decrease in phosphorylated RPS6 signal in all mitotic-enriched samples; however, this is not significant relative to asynchronous levels of phosphorylated RPS6 by one-way ANOVA (Figure 3D, E). Previous research suggests that the hyperphosphorylation of 4E-BP1 in mitosis is independent of the mTOR pathway (32). Collectively, our results demonstrate that 4E-BP1-dependent translation inhibition is not a mechanism that reduces protein synthesis during mitosis. Coupled with our analysis of ISR-dependent inhibition of translation, these results suggest that inhibition of translation initiation is not solely responsible for mitosis-associated inhibition of protein synthesis.

### Elongation rate is slowed in a prolonged mitosis

After investigating mitotic impact on initiation, we sought to determine whether elongation was affected. In stress, eEF2 is phosphorylated by eEF2 kinase (eEF2K) to inhibit translation elongation (40). We used phosphate buffered saline (PBS) treatment as a control which simulates amino acid starvation and triggers eEF2 phosphorylation. Interestingly, we also observed increased phosphorylation of eEF2 in both cycling and prolonged mitoses (Figure 4A, B). This suggests there is decreased translational elongation in mitosis because eEF2 translocates the growing polypeptide from the aminoacyl (A-)site to the peptidyl (P-)site in the 80S ribosome, allowing for further amino acid deposition into the now-empty A-site (41). Phosphorylation of eEF2 renders it inactive, suggesting limited translocation in mitosis.

**Figure 4.**
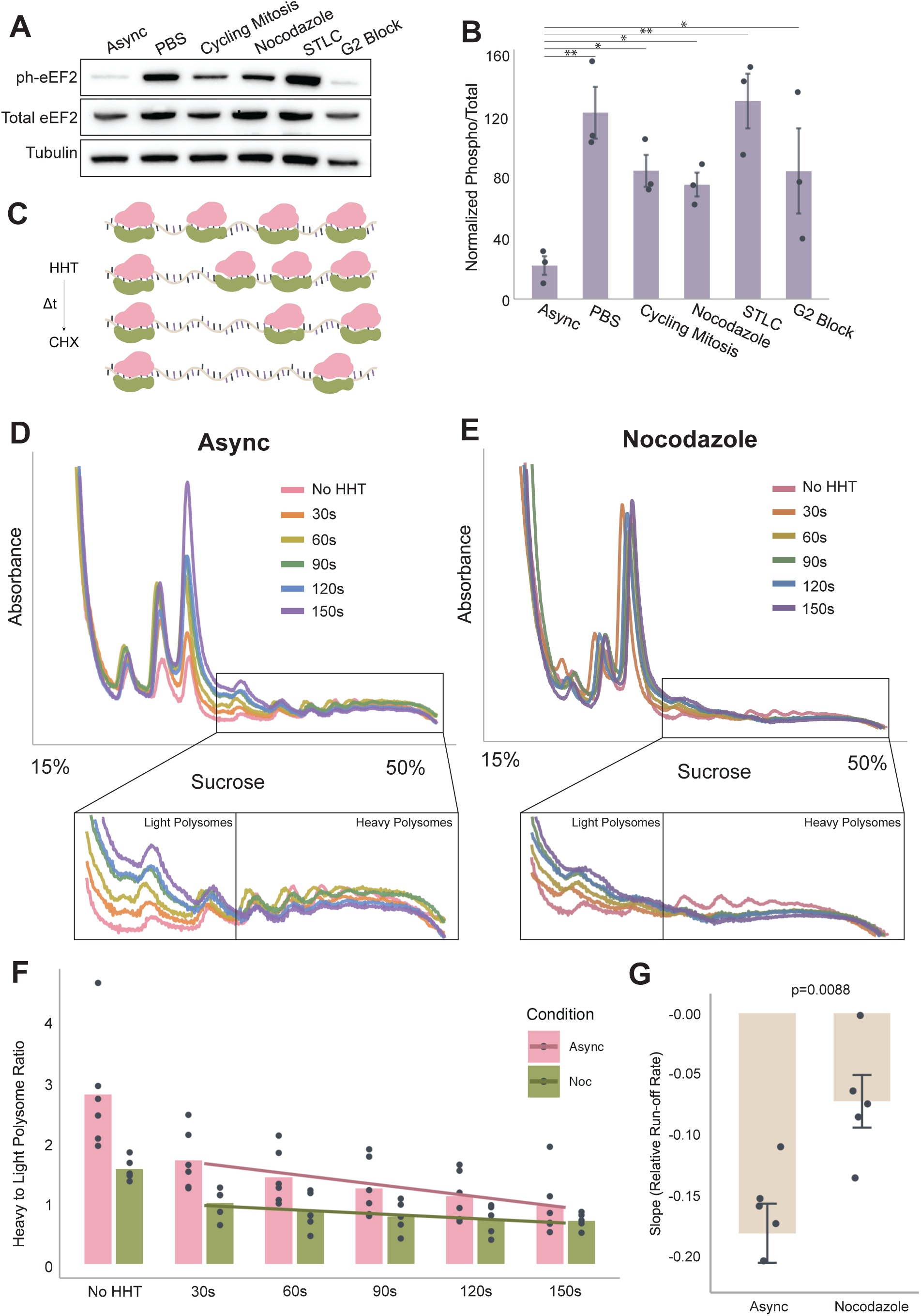
eEF2 is phosphorylated and elongation rate is slower in mitosis. **(A)** Western blot of phosphorylated and total eEF2 in RPE-1 cells. **(B)** Quantification of normalized phosphorylated to total eEF2 signal intensity. Statistics determined by one-way ANOVA with Dunnett’s post-hoc test, asynchronous as reference. Error bars represent the standard deviation, n=3 biological replicates. **(C)** Method schematic of HHT addition over time and ribosomal run-off on transcripts. **(D, E)** Polysome traces of asynchronous **(D)** and mitotic **(E)** RPE-1 cells in HHT for 30-150s; main traces shown in full, with a zoomed-in panel focusing on the polysome region (below). **(F)** Ratio of the area under the curve of the heavy (4+ ribosomes) to light (2-3 ribosomes) polysomes over time in HHT. Line fit to 30-150s HHT. Slope of line quantified in **(G)**. Significance determined by Student’s T-test, error bars calculated by standard error of the mean, n=5 biological replicates.

To directly test whether the elongation rate was slowed in a prolonged mitosis, we treated asynchronous and nocodazole-treated cells with homoharringtonine (HHT) in a time course, followed by cycloheximide (CHX) treatment to study the rate at which ribosomes run off (Figure 4C). These methods are similar to those described in previous literature (42). After HHT-treating asynchronous and prolonged mitotic cells for 0-150 seconds at 30 second intervals, we analyzed lysates through polysome profiling. Our analysis shows a gradual increase in the area under the monosome peak in asynchronous cells from 0-150 seconds, suggesting an increase in the number of transcripts with the single initiating ribosome remaining. Additionally, we see a gradual increase in the number of transcripts with two or three ribosomes (termed here as “light polysomes”), with a concurrent decrease in the number of transcripts with four or more ribosomes (“heavy polysomes”) (Figure 4D). We recorded a similar trend in ribosomal run-off with our nocodazole-treated mitotic cells, but the difference between timepoints was much less robust (Figure 4E).

To evaluate the difference in the relative run-off rate between asynchronous and cells in a prolonged mitosis, we compared the area under the heavy polysomes by the area under the light polysomes for each HHT treatment to determine the change in the number of transcripts with fewer ribosomes over time (Figure 4F). The slope of the line fit to this quantitation reveals that the relative run-off rate in a prolonged mitosis is slower than that of its asynchronous counterpart (Figure 4G). This difference in slope suggests that ribosomes are transversing mRNA more slowly, further indicating the elongation rate is slowed in a prolonged mitosis.

### tRNA-Alanine-CGC availability is decreased in mitosis

Previous literature suggests fewer tRNAs are associated with polysomes in mitosis (43). We hypothesized that the phosphorylation of eEF2 could be triggered by a decrease in the availability of cognate-tRNAs leading to an empty A-site (44, 45). In the event that a cognate-tRNA is unavailable for deposition of its amino acid, this would cause stalling of the ribosome at that particular codon (46). If specific tRNAs are depleted during mitosis, we expect to see an increase in A-site occupancy of particular codons with low cognate-tRNA abundance in mitosis. If all tRNAs are decreased during mitosis, we do not expect to observe an increase in A-site occupancy of any codons. To evaluate this, we reanalyzed ribosome profiling data from interphase, cycling, and prolonged mitosis, to determine the frequency at which codons occupy the A-site in mitosis (47). We aligned ribosomal footprints to cDNA and counted the frequency of each codon in the A-site position for each dataset, normalized to total codon frequency across ribosomal associated sequences. We compared codon frequencies between interphase and cycling mitosis (Figure 5A, upper panel) and prolonged mitosis (lower panel). Positive values have higher A-site occupancy in interphase; negative values have higher A-site occupancy in mitosis. Interestingly, the three stop codons, UAA, UGA, UAG, have a higher occupancy in interphase relative to a prolonged mitosis (Figure 5A, lower panel), which could suggest that elongation is slowed such that ribosomes do not reach the end of the opening reading frame at the same frequency as asynchronous cells. This is also supported by our relative runoff rate data (Figure 4). Furthermore, both cycling and prolonged mitotic footprints indicated a higher frequency of the codon Alanine-GCG in the A-site relative to interphase cells (Figure 5A, arrow).

**Figure 5.**
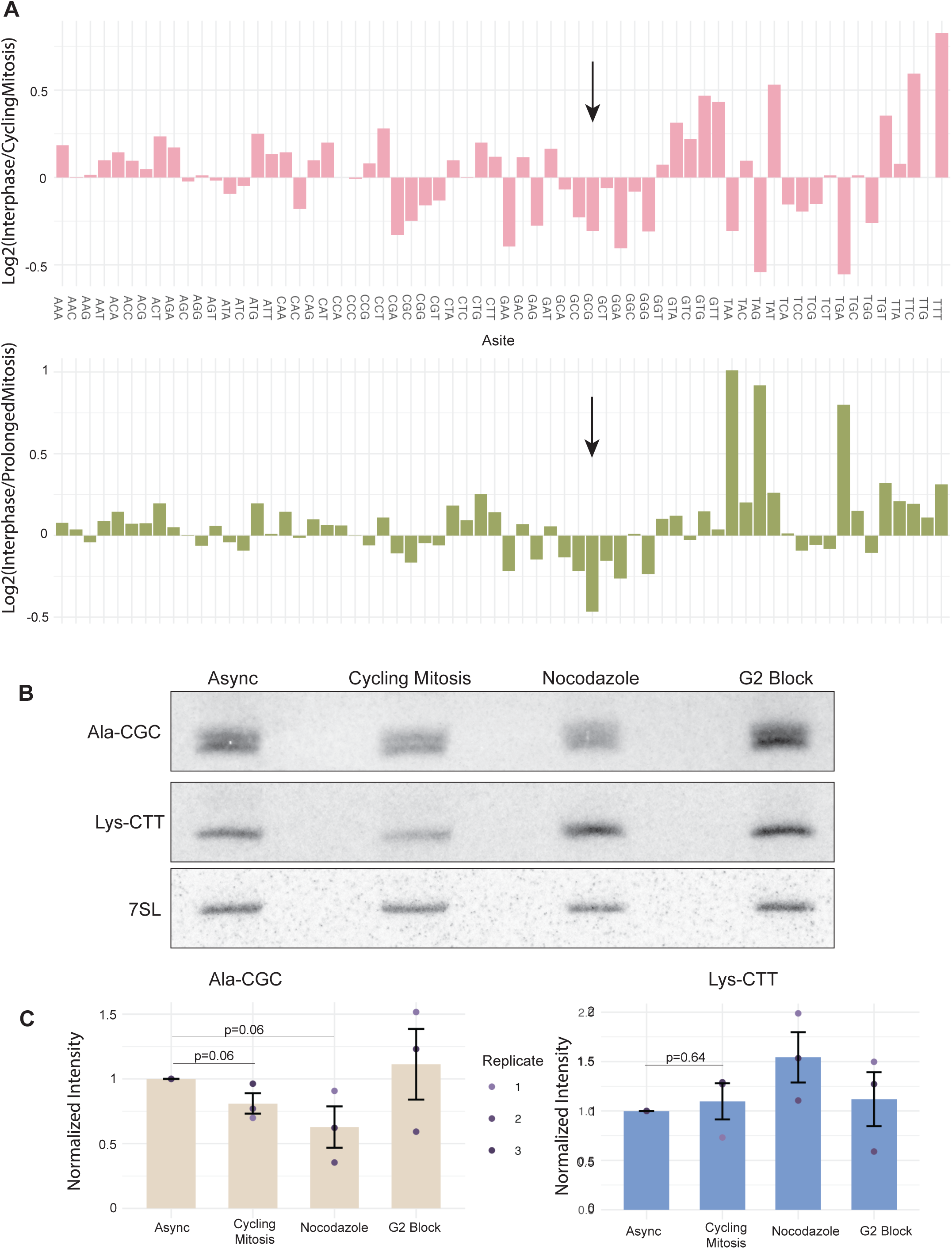
tRNA-Alanine-CGC availability is decreased in mitosis. **(A)** Codon usage plots in cycling (top) and prolonged (bottom) mitotic HeLa cells (47). **(B)** Northern blot of tRNA-Ala^CGC^ and tRNA-Lys^CTT^ tRNAs. **(C)** Quantification of northern blot analysis. Statistical significance determined by Wilcoxon rank sum test. Error bars represent the standard deviation, n=3 biological replicates.

This increased occupancy in the A-site during mitosis may suggest a decreased availability of its cognate-tRNA, tRNA-Ala^CGC^, in mitosis. To test this, we assayed total tRNA levels in asynchronous and mitotic samples through northern blotting. This revealed a decrease in total tRNA-Ala^CGC^ in mitosis, with a trend toward fewer available in a longer mitosis (Figure 5B, C, left panel). We compared this to tRNA-Lys^CTT^ which had no difference in codon occupancy (AAG) in mitosis relative to interphase through bioinformatic analysis and subsequently had no significant decrease in total levels in mitosis (Figure 5B, C, right panel). Targeted reduction of a single tRNA transcript level has recently been observed in response to DNA damage, which has a similar single-isoacceptor effect by tRNA-Leu^UAA^ (30).

### Inhibition of RNAPolIII contributes to a decrease of translation

Decreased tRNA abundance could be caused by increased degradation rate or a decreased synthesis rate. Mitotic transcriptional inhibition (4, 48) will lead to a significant decrease in tRNA synthesis that will be exacerbated by increased time in mitosis. Production of tRNAs is mediated by RNAPol III, a polymerase also affected by global transcriptional shutdown in mitosis (5). To evaluate whether a lack of tRNA synthesis phenocopies decreased translational elongation, we examined phosphorylation of eEF2 and eIF2α with RNAPol III inhibitor-treated asynchronous cells. We found eEF2 exhibits significantly increased phosphorylation following inhibition of RNAPol III in asynchronous cells; however eIF2α was not phosphorylated relative to asynchronous cells (Figure 6A-D). This result is similar to that of PBS treatment, in which starvation leads to phosphorylation of eEF2, but not eIF2α (Figure 6A-D). Further, polysome profiling following RNAPol III inhibition indicates that loss of tRNA transcription contributes to an attenuation of translation, although not to the extent of a prolonged mitosis (Figure 6E). Therefore, we conclude that the natural inhibition of Pol III in mitosis, preventing further synthesis of tRNAs, contributes to the decrease in translation in mitosis.

**Figure 6.**
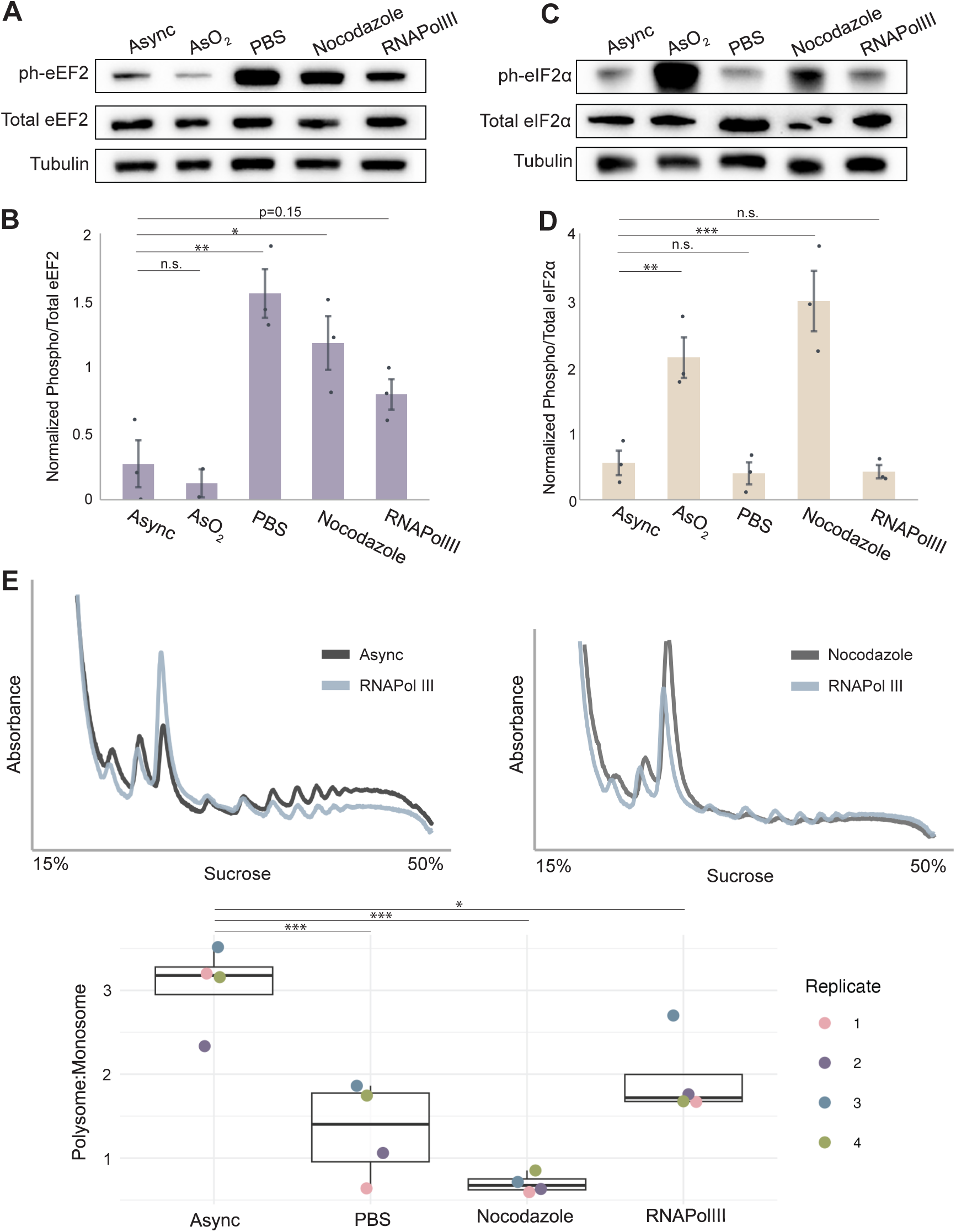
RNAPol III inhibitor phosphorylates eEF2 and contributes to decreased translation efficiency. **(A)** Western blot of phosphorylated and total eEF2 in RPE-1 cells. **(B)** Quantification of normalized intensity of phosphorylated eEF2 to total eEF2. Statistical significance determined by one-way ANOVA with Dunnett’s post-hoc test, asynchronous as reference. Error bars represent the standard deviation, n=3 biological replicates. **(C)** Western blot of phosphorylated and total eIF2α in RPE-1 cells. **(D)** Quantification of normalized intensity of phosphorylated to total eIF2α, significance determined by one-way ANOVA with Dunnett’s post-hoc test. **(E)** Polysome traces of asynchronous and Pol III inhibited cells (left) and nocodazole-treated and Pol III inhibited (right) RPE-1 cells. Quantified (below) by polysome to monosome ratio to determine translation efficiency. Statistics determined by one-way ANOVA with Dunnett’s post-hoc test.

## Discussion

Our work evaluates the effects of mitosis on translation by multiple enrichment strategies (Supplemental Figure 1), observing a consistent decrease in global translation (4, 6) through polysome profiling, metabolic labeling, and puromycin labeling (Figure 1, Supplemental Figure 2). The novelty of this work focuses on a primary, immortalized cell line, immortalized hTERT-RPE-1 cells, which have not adapted alternative mechanisms of cell division. We find there are multiple mechanisms by which protein synthesis decreases in mitosis, with a trend toward increased inhibition with prolonged time in mitosis. We find elongation rate is regulated in mitosis, coincident with a decreased abundance of tRNA-Ala^CGC^, which we hypothesize causes ribosomal pausing. We speculate that the inability to replenish the availability of this tRNA is due to the natural inhibition to Pol III transcription in mitosis.

We have shown that the effect of the translational attenuation is, in part, mediated by regulation at initiation. Cycling mitotic cells show a slight increase in phosphorylated eIF2α, which increases the longer cells are in mitosis, such as those synchronized in nocodazole and STLC (Figure 2A, B). However, chemical or genetic bypass of the effects of eIF2α phosphorylation on translation did not restore mitotic translation to levels observed in interphase (Figure 2C-F). This suggested that multiple mechanisms repress protein synthesis during mitosis. Further work is necessary to discern which kinase is responsible for phosphorylating eIF2α in mitosis. We hypothesize that GCN2 is the active kinase during mitosis since it responds to amino acid starvation (49). Additionally, we find that 4E-BP1 is unlikely to affect the formation of eIF4F on the cap of mRNA, as it does not exhibit increased binding to eIF4E1 during mitosis (Figure 3A-C). Mitosis may slightly suppress the mTOR pathway, but we do not believe it drives translational repression in mitosis (Figure 3D, E).

In addition to suppression of translation initiation, we confirm phosphorylation of eEF2 in mitosis (Figure 4A-B, (23)), suggesting that mitotic translation is also suppressed through an effect on elongation rate (50). In support of this hypothesis, we found that cells in a prolonged mitotic arrest show a slower ribosomal run-off rate relative to asynchronous cells (Figure 4), confirming that elongation rate in mitosis is decreased. Combined with our data revealing mitotic suppression of translation initiation, we conclude translation is regulated at multiple stages in mitosis.

Phosphorylation of eEF2 occurs upon amino acid starvation (Figure 4A, (51)). Previous literature suggested fewer tRNAs associated with polysomes in mitosis (43) suggesting either a lack of tRNA abundance or a decrease in charged tRNA available. To determine whether a depletion of specific tRNAs triggered eEF2 phosphorylation, we compared the occupancy of codons in the A-site in ribosome profiling data for mitotic and interphase cells (47). The increased occupancy of the Alanine-GCG codon in the A-site of mitotic ribosomes (Figure 5A) indicated potential stalling at this codon due to a decreased availability of its cognate tRNA, tRNA-Ala^CGC^. Northern blotting confirmed a specific decrease in total tRNA availability of tRNA-Ala^CGC^. In contrast, Lysine-AAG codon occupancy in the A-site was similar in asynchronous and mitotic libraries, and its cognate tRNA abundance, tRNA-Lys^CTT^, did not change (Figure 5B, C).

Because the half-lives of tRNAs are approximately 50 hours in eukaryotic cells (52), it is unlikely that the intrinsic turnover rate of Ala^CGC^ is responsible for its decreased abundance in both a cycling and prolonged mitosis. Since mitosis persists for about an hour in RPE-1 cells (53), it is more likely that this change in abundance is caused by increased tRNA degradation during mitosis. Previous literature has indicated tRNA abundance is dynamic in response to different environmental conditions. For example, modifications to tRNAs in cancer can increase abundance of certain tRNAs, here Arg-TCT-4-1, which leads to translation of specific ACA-codon-heavy transcripts to promote oncogenesis (29). Additionally, during DNA damage, research has shown that a decrease in tRNA-Leu^UAA^ causes ribosome stalling as a p53-independent mechanism of apoptosis (30). This recent literature suggests that cells alter the availability of specific tRNAs to promote cell growth and respond to stress. We propose that decreased tRNA-Ala^CGC^ is a component of a general pathway that decreases translation during mitosis.

Further research needs to be conducted to understand the mechanisms by which this specific tRNA is degraded in mitosis. However, since RNAPolIII activity is inhibited naturally in mitosis (5), we believe the failure to replenish the tRNA-Ala^CGC^ in mitosis contributes to the regulation of translation in mitosis. The inability to maintain basal levels of tRNA-Ala^CGC^ is likely to trigger the phosphorylation of eEF2, eIF2α, and decrease the rate of translation in mitosis. To explore this, we treated asynchronous cells with a RNAPolIII inhibitor, which revealed a modest increase in phosphorylation eEF2, indicating regulation at the elongation step (Figure 6A). However, inhibition of RNAPolIII did not lead to phosphorylation of eIF2α (Figure 6B). Modulation of elongation does have a modest effect on translation efficiency with Pol III inhibition (Figure 6C), which suggests that the natural inhibition of transcription in mitosis contributes to a decreased elongation rate in a prolonged mitosis. Our data suggest that tRNA-Ala^CGC^ decreased abundance, and failure to replenish working levels due to repressed transcription in mitosis, triggers eEF2 phosphorylation to regulate translation elongation. This is a starvation response which causes eIF2α phosphorylation to regulate initiation, preventing ribosomal collisions.

We further speculate that this mechanism of tRNA dynamics is a regulatory response for cell cycle progression. In bacteria, a similar phenomenon has been observed in response to contact-dependent growth inhibition, in which regulation of cell growth activates a typically-latent RNase to target tRNAs. Other toxin-antitoxin responses in bacteria have been shown to target specific tRNAs as a mechanism against bacteriophage infection or toward bacterial competition (54, 55) Recent literature in eukaryotes has reported an attenuation of protein synthesis during mitosis as quality control for cell proliferation, triggering a p53-dependent cell cycle arrest in cells that have been arrested in mitosis for an extended time (53). Regulation of tRNA abundance is observed in both prokaryotes and eukaryotes. Therefore, the targeted loss of certain tRNAs, with an enhanced effect the longer cells are in mitosis, may function similarly to regulate the translation of alanine-rich transcripts. We speculate that the downregulation of tRNA-Ala^CGC^ regulates cell proliferation, as a subset of transcripts related to the ERK pathway are alanine-rich (56). The ERK pathway has implications in cell proliferation (57), so perhaps the slowed translation of ERK-related proteins is a regulatory step to dampen MAPK signaling as the cell actively divides in mitosis, resuming after the completion of this cell cycle. Further research should explore the effects on translation and cell cycle progression through restoring tRNA availability in mitotic cells.

## Acknowledgements

The authors thank Daniel Cifuentes for use of the Amersham Typhoon Phosphor Imager. We acknowledge that much of the computational work reported in this paper was performed on the Shared Computing Cluster which is administered by Boston University’s Research Computing Services. This work was supported by grants from NIGMS to M.B. (5R01GM144352-03, 5R01GM122893-07)

---

**Supplemental Figure 1.**
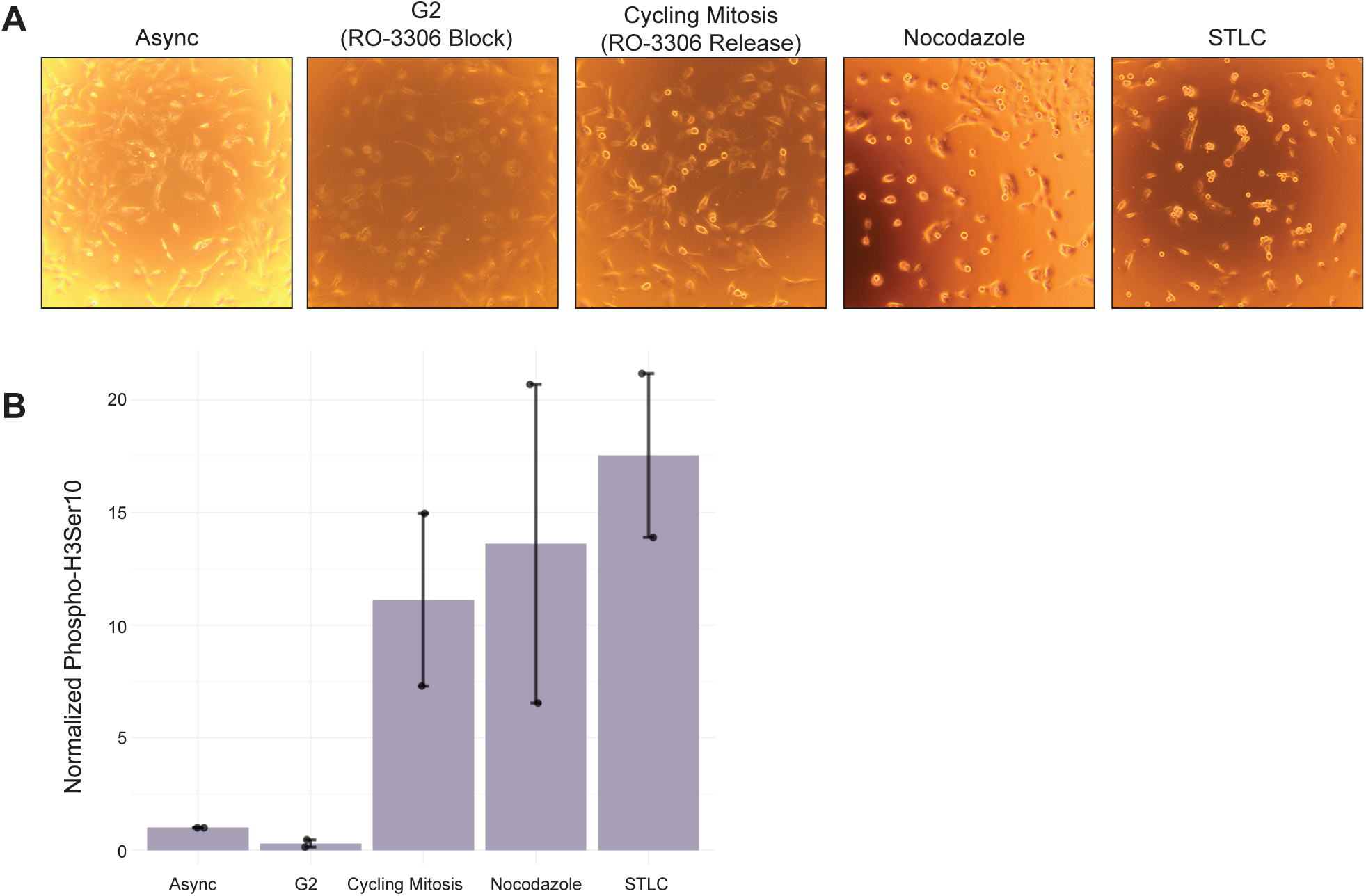
Confirmation of drug treatment for mitotic enrichment. **(A)** Phase images of drug synchronizations for mitotic enrichment. Asynchronous cells consist of no drug treatment; cycling mitosis yields mitotic enrichment 45 minutes after release from RO-3306; mitotic enrichment with nocodazole for 16 hours; mitotic enrichment with STLC for 16 hours; G2 enriched cells blocked in RO-3306 for 16 hours (left to right). Rounded cells are mitotic. **(B)** Western blot quantification of Phospho-Histone H3Ser10 for mitotic enrichment on 15% SDS-PAGE gel.

**Supplemental Figure 2.**
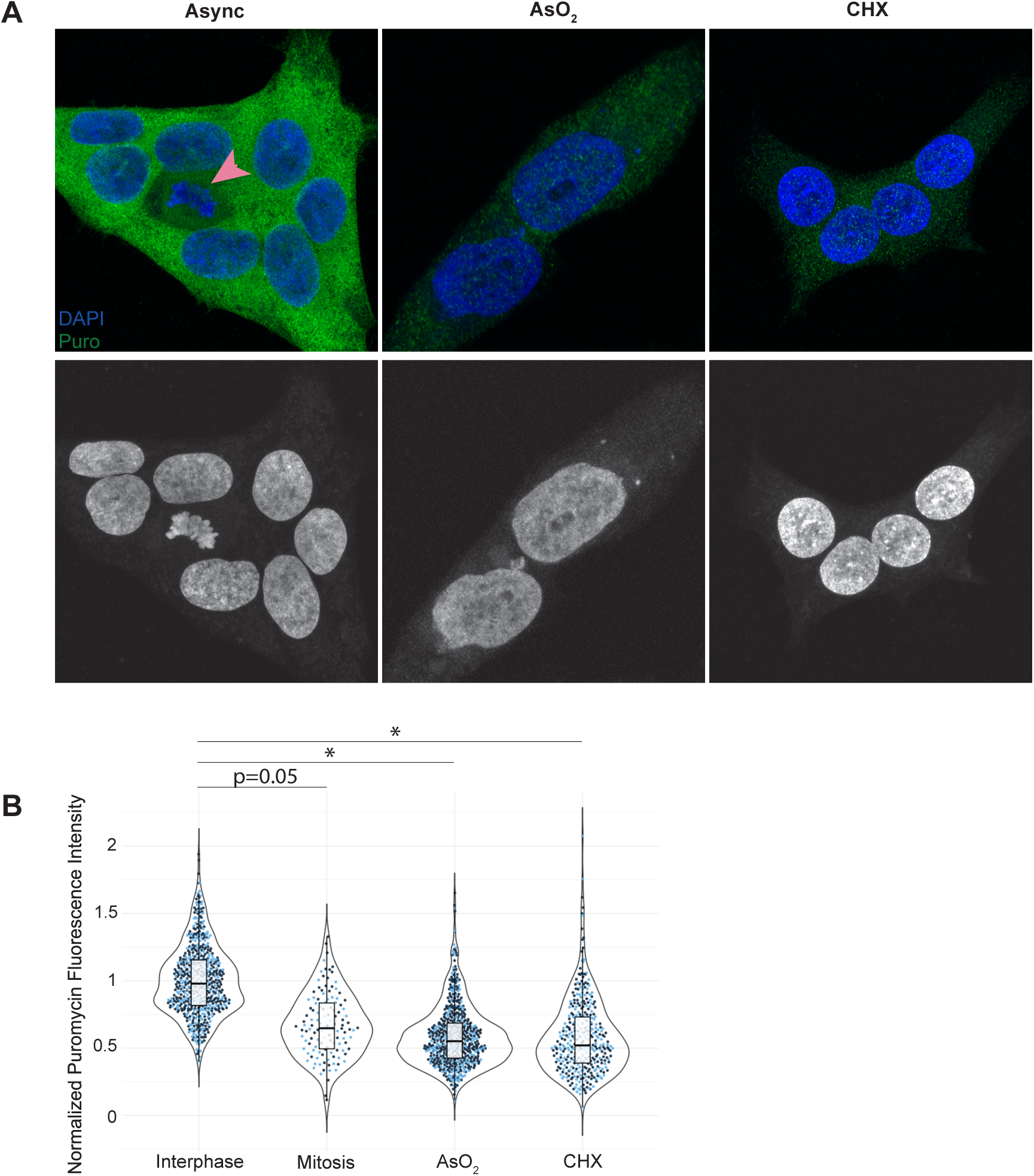
Basal translation of mitotic cells decreased in asynchronous population. **(A)** Immunofluorescence of puromycin in asynchronous population of Hap1 cells, sodium arsenite treated (AsO_2_), and cycloheximide (CHX) treated. Mitotic cell indicated by pink arrow. **(B)** Quantification of normalized mean fluorescence intensity, relative to interphase mean. Significance determined by Student’s paired T-test (biological replicates, n=2).

**Supplemental Figure 3.**
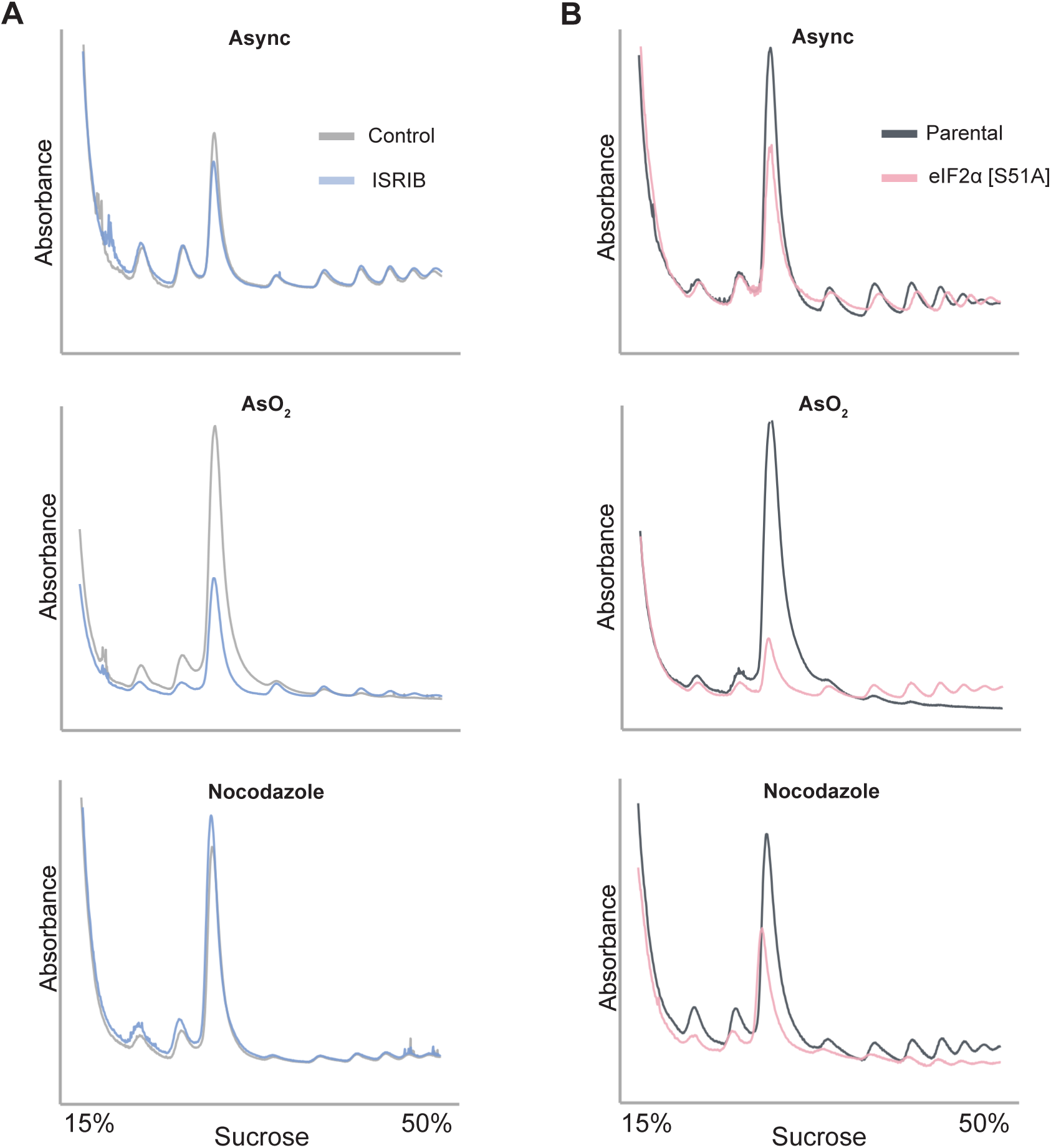
Polysome traces for RPE-1 ISRIB treatment and Hap1 eIF2α [S51A] cells. **(A)** RPE-1 co-treatment of sodium arsenite and ISRIB or Nocodazole and ISRIB polysome traces. **(B)** Hap1 eIF2α [S51A] cells treated with Nocodazole compared to parental.

**Supplemental Figure 4.**
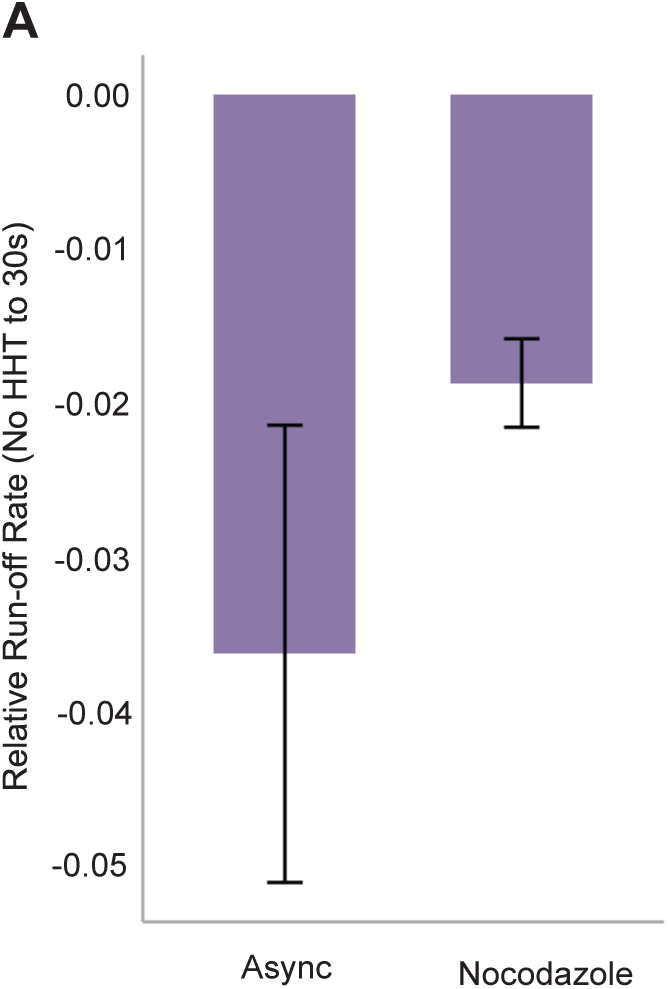
Relative ribosomal run-off rate after 30 seconds of HHT treatment. **(A)** Slope of asynchronous and mitotic cells between no HHT treatment and 30 seconds of HHT treatment. Showing n=5 biological replicates, error bars represent standard deviation.

## Materials and Methods

### Cell culture, cell synchronization, and drug treatments

hTERT-immortalized RPE-1 cells were cultured in DMEM-F12 supplemented with 10% heat-inactivated fetal bovine serum, 1 U ml^-1^ penicillin-streptomycin, and sodium bicarbonate at 37°C with 5% CO_2_. Cell lines routinely tested for Mycoplasma contamination. Hap1 cells were cultured in DMEM media supplemented with 10% heat-inactivated fetal bovine serum and 1 U ml^-1^.

Asynchronous cells are untreated and adherent cells collected. For cycling mitotic cells, RPE-1 cells are treated at 60% confluency with RO-3306 (Selleckchem, S7747) at 10 μM final concentration for 16 hours. Cells are washed 5 times in PBS and released into drug-free media for 45 minutes. Miotic-enriched cells are collected by mitotic shake off. Prolonged mitotic cells are treated with nocodazole (100nM, Selleckchem, S2775) or STLC ((+)-S-Trityl-L-cysteine) (50μM, Sigma 164739) at 60% confluency for 16 hours and collected by mitotic shake off. G2 synchronized cells are treated with RO-3306 for 16 hours and adherent cells are collected. Sodium arsenite treated asynchronous cells were subjected to 250-500μM sodium arsenite (Sigma S7400) for an hour. PBS treated asynchronous cells were subjected to 1X PBS for 1 hour. RPE-1 cells treated with ISRIB (5 μM) were either treated for one hour in asynchronous cells, co-treated with sodium arsenite (500μM) and ISRIB for an hour, or nocodazole (100nM) and ISRIB for 16 hours. An mTOR1/2 inhibitor (INK-128, HY-13328, MedChemExpress) was added to asynchronous cells at 200nM for one hour. Inhibition of Pol III was mediated by ML-60218 (MedChemExpress), through overnight (16h) treatment of asynchronous cells at 90μM.

### Polysome analysis

Basal polysome profiles were performed according to He and Green (58) with modifications. Cells were incubated with 100 μg mL^-1^ of cycloheximide (Fisher, #357420010) for 10 minutes, then mitotic cells were shaken off, spun down, and harvested and asynchronous and G2-blocked cells were harvested in lysis buffer (10mM HEPES-KOH, pH 7.4, 150mM KCl, 10mM MgCl_2,_ 1mM DTT, 100ug mL^-1^, 2% NP-40, HALT Protease inhibitor, 6U mL^-1^ Promega RNase Inhibitor). Cells were lysed for 5 minutes on ice; lysate was cleared by centrifugation at 17, 000xg for 5 minutes at 4°C. Sample was quantified and 1-1.2 μg protein run on a 15-50% sucrose (10mM HEPES-KOH [pH 7.4], 150 mM KCl, 10 mM MgCl_2_, 1 mM DTT, and 100 μg mL^-1^ CHX) gradient using Beckmann Coulter SW41 Ti rotor at 40, 000 rpm at 4°C for 2 hours. Sucrose is supplemented with Halt Protease Inhibitor Cocktail (Thermo Fisher, 78430). Gradients are run on BioComp Gradient Master reading A260. Samples treated with homoharringtonine (Sigma, ML1091) at a final concentration of 3.7 μM for 30-150 seconds at 30 second intervals, followed by CHX treatment.

### Metabolic Labeling

Cells were cultured in methionine-free media supplemented with respective drug treatments for 15 minutes, followed by labeling with [^35^S]-methionine/cystine for 15 minutes to assess global protein synthesis. After washing and lysis, 10-15 μg total protein was resolved on TGX Stain-Free FastCast Acrylamide 12% gels (Bio-Rad, #1610185), followed by transfer by Bio-Rad Trans-blot Turbo Transfer system. Dry membranes were exposed to a phosphor screen overnight, and imaged by the Cytiva Amersham Typhoon Biomolecular Imager. Quantification was analyzed by BioRad Image Lab Software.

### Puromycin Immunofluorescence

Cells were subjected to 250 μM sodium arsenite (Sigma S7400) for 50 minutes or 100 μg mL^-1^ of cycloheximide (Fisher, #357420010) for 10 minutes, or drug-free media for asynchronous cells. A final concentration of 10 μg mL^-1^ of puromycin was added for 10 minutes followed by 4% PFA fixation for 10 minutes. Cells were permeabilized with 0.5% Triton-X for 15 minutes. Coverslips were incubated with Puromycin (see Antibodies) at 1:200 concentration overnight at 4°C, followed by secondary incubation with goat anti-mouse AlexaFluor-488 (708-545-149, Jackson Immunoresearch) for 1 hour at 37°C. Images taken on Nikon A1R Confocal Microscope. Quantification was analyzed by ImageJ.

### Western blot analysis

Cells were lysed in NP-40 Lysis Buffer (150mM NaCl, 1% NP-40, 50mM Tris [pH 8.8], Halt Protease inhibitor). Lysate was cleared and quantified by Bradford assay. A total of 10-15ug of protein was separated by TGX Stain-Free FastCast Acrylamide (#1610185) 12% gel electrophoresis (or sodium dodecyl sulfate-15% polyacrylamide gel for 4E-BP, Figure 3). Proteins were transferred to a nitrocellulose membrane (Cytiva, Amersham) by Bio-Rad Trans-blot Turbo Transfer system. Blots were incubated in 5% PBS-T milk, followed by primary incubation overnight. After washing, blots were incubated in anti-host-HRP secondary for 1 hour at room temperature, followed by chemiluminescent exposure by the Bio-Rad ChemiDoc. Quantification was analyzed by ImageJ.

### Antibodies

Phospho-eIF2α - Abcam ab32157, Rabbit

Total-eIF2α - Cell Signaling Technology #9722, Mouse

Total 4E-BP1 (53H11) – Cell Signaling Technology #9644, Rabbit

Non-phospho-4E-BP1 (Thr46) – Cell Signaling Technology #4923, Rabbit

Phospho-4E-BP1 (Ser65) – Cell Signaling Technology #9456, Rabbit

Phospho-4E-BP1 (Thr70) – Cell Signaling Technology #13396, Rabbit

eIF4E – Cell Signaling Technology #9742, Rabbit

Total eEF2 – Cell Signaling Technology #2332, Rabbit

Phospho-eEF2 Thr56 – Cell Signaling Technology #2331, Rabbit

DM-1A Tubulin – Sigma #T9026, Mouse

GAPDH – Santa Cruz #sc-25778, Rabbit

Phospho-Histone H3Ser10 – Proteintech #66863-1-Ig, Mouse

Puromycin – Millipore Sigma #MABE343, Mouse

### Northern Blot Analysis

Total RNA extraction was performed using TRIzol reagent (Thermo Invitrogen, 15596026), following associated protocol. Between 10 and 20 μg of total RNA was loaded on a 12% TBE-urea gel in 2X Urea RNA Loading dye (see methods, (59)), boiled at 85°C for 5 minutes, prior to loading. Samples were run for 2-3 hours at 300 V. Gels were SYBR gold stained for 10 minutes and transferred to a positively charged nylon membrane (GE Healthcare, RPN303B) for 45 minutes at 300mA using the BioRad Trans-Blot Turbo System. The membrane was air dried for an hour prior to UV crosslinking. Blots were pre-hybridized in blocking (5X SSC, 5X Denhardt’s solution, 0.5% SDS, 25 μg/mL denatured salmon sperm DNA) for an hour. Blots were then hybridized with ^32^P probe in ULTRAhyb Ultrasensitive Hybridization buffer (Thermo Invitrogen, AM8670). Probes include tRNA-Alanine-CGC (5’-GCGCTCTACCACTGAGCTACA-3’), tRNA-Lysine-CTT (5’-TACCGACTGAGCTAGCCGGGC-3’), 7SL (5’- GCTCCGTTTCCGACCTGGGCC-3’), prepared with 50 μCi ^32^P. After washes (2X SSC, 0.1% SDS), membrane was exposed to phosphor screen for 10-16 hours and imaged on the Cytiva Amersham Typhoon Biomolecular Imager. Quantification of tRNA signal was normalized to 7SL signal.

### Cap Pulldown Assay

Cells synchronized as previously described were lysed in NP-40 Lysis Buffer (see *Western blot analysis*). Jena Bioscience m^7^GTP-sepharose beads (NU-1122) were incubated with 600-1000 μg of protein and tumbled for one hour. Beads were washed twice and eluted by centrifugation. Eluates were subsequently run on a gel as described in Western blot analysis.

### Codon usage plot analysis

Data was downloaded from Gene Expression Omnibus (GSE230189). Ribosome profiling data was processed by trimming adapters with CutAdapt using the following paramaters: cutadapt -m 15 -u 8 -e 0.1 --match-read-wildcards -a TCGTATGCCGTCTTCTGCTTG -O 1 -o. Trimmed reads were aligned to human rRNA sequences and aligning reads were removed: -q $fq --sam --seedmms 2 --seedlen 11 --seed 494123 --maqerr 70 --tryhard -k 1 --un $out --best --maxbts 800. Non-rRNA reads were then aligned to the human Refseq Select cDNA sequences using Bowtie with the following paramaters: --sam --seedmms 2 --seedlen 11 --seed 494123 --maqerr 70 --tryhard -k 1 --best -- maxbts 800. Header information was then removed form each .sam file using samtools. To determine the P-site offset for each library we read each aligned library into R and identified reads that intersect with the annotated start codon. We then calculated the distance from the start codon to the 5’ end of the read (P-site offset) using a custom R-script (OffsetPlots_Batch.Rmd). From this analysis we determined the most common offset for both the P-site and A-site. We then used the calculated P- and A-site offsets to measure codon frequency in each site. We read aligned libraries into R, selected reads of length 32 or 33nt and counted the frequency of each in-frame codon in the P-site and A-site using a custom R script (Offset13Batch.Rmd). To normalize for differences in codon content in different libraries we calculated codon frequencies across the entire length of each ribosome-associated read for each aligned library. We read in aligned libraries to R, determined the reading frame of each read based on the distance from the start codon and calculated the frequency of all codons across all reads using a custom R script (Codons.Rmd). To generate codon frequency normalization factors we calculated ratios of codon percentages for pairs of libraries (e.g. IF1 vs cycling M1, etc). We then applied these scaling factors to comparisons of codon percentages calculated for A-sites.

